# TMS-EEG reveals causal dynamics of the premotor cortex during musical improvisation

**DOI:** 10.1101/2025.10.04.680222

**Authors:** Pedro T. Palhares, Sasha D’Ambrosio, Óscar F. Gonçalves

## Abstract

Musical improvisation illustrates the brain’s capacity for flexible, creative motor control, yet the causal mechanisms underlying this complex behaviour remain poorly understood. We employed transcranial magnetic stimulation combined with electroencephalography (TMS-EEG) to probe state-dependent cortical dynamics in the left dorsal premotor cortex (PMd) of professional jazz pianists (n = 3) during improvisation, sight-reading, and rest. This proof-of-concept study demonstrates the feasibility of combining perturbational neuroscience with ecologically valid musical performance. Multiple convergent analyses revealed distinct cortical signatures during improvisation: reduced local mean field power, decreased phase-locking of evoked responses, and preserved but gain-modulated early components as revealed by Correlated Components Analysis. These findings suggest that improvisation is characterized by attenuated PMd excitability and more variable response timings, while preserving the fundamental architecture of cortical responses. This perturbational signature supports a neural efficiency model of expertise whereby expert musicians achieve creative flexibility through training-induced streamlined, optimized cortical processing. Our results establish TMS-EEG as a powerful approach for investigating the causal dynamics of creative cognition and demonstrate how the brain reconfigures its response properties to support internally driven motor performance.

## Introduction

Musical performance represents one of the most sophisticated forms of human sensorimotor behaviour, requiring the precise coordination of perceptual, cognitive, and motor systems in real time (Vuust et al., 2022; Zatorre et al., 2007). Within this domain, musical improvisation offers a particularly compelling model for understanding flexible motor control and creative cognition (Palhares et al., 2024), demanding the spontaneous generation of novel motor sequences while maintaining temporal precision and adhering to complex harmonic constraints (Pressing, 1988). Despite extensive neuroimaging research on the neural correlates of musical improvisation (Beaty, 2015; Loui, 2018), the causal dynamics underlying this complex behaviour remain poorly understood, particularly regarding the functional role of premotor regions in orchestrating improvised (internally generated) versus externally cued motor performance.

The premotor cortex (PMC) plays a critical role in motor planning, sequence generation, and the transformation of abstract goals into concrete motor commands (Hoshi & Tanji, 2007; Rizzolatti et al., 1996). In the context of musical improvisation, neuroimaging studies have shown that the PMC undergoes functional reorganization, displaying altered connectivity patterns with other nodes of the motor network, including increased coupling with prefrontal control regions and sensorimotor areas (Donnay et al., 2014; Pinho et al., 2014). Improvisation thus appears to engage a complex interplay between idea generation and evaluation processes, requiring the coordination of multiple brain networks which include auditory-motor regions for perception-action cycles and both medial and lateral prefrontal areas for creative cognition (Beaty, 2015; Loui, 2018; Pinho et al., 2015). However, the correlational nature of neuroimaging data precludes strong causal inferences about how the PMC actively contributes to these distinct performance states (Havlík et al., 2023; Logothetis, 2008; Miniussi & Thut, 2010).

Transcranial magnetic stimulation combined with electroencephalography (TMS-EEG) offers a unique window into the causal dynamics of cortical function by directly perturbing neural activity and measuring the resulting spatiotemporal response patterns (Ilmoniemi et al., 1997; Miniussi & Thut, 2010). Unlike traditional neuroimaging approaches that measure correlates of neural activity, TMS-EEG provides direct access to cortical excitability, connectivity, and state-dependent processing through the analysis of TMS-evoked potentials (TEPs). The morphology, amplitude, and spatiotemporal spread of TEPs reflect intrinsic properties of the stimulated cortical region and its connected networks, with evidence suggesting that these responses are sensitive to ongoing behavioural states (Nikulin et al., 2003) and level of consciousness (Sarasso et al., 2015).

The distinction between improvised and sight-read musical performance in contrast with task-free resting state provides an ideal paradigm for investigating task-dependent variations in cortical responses to perturbation. Improvisation requires the integration of multiple sources of information — harmonic knowledge, motor memories, auditory feedback — into a coherent but flexible motor output. In contrast, sight-reading involves a more constrained transformation from visual input to motor output. How these different task demands are reflected in the brain’s response to direct perturbation remains an open question.

In this study, we employed TMS-EEG to directly probe the state-dependent dynamics of the left dorsal premotor cortex during musical improvisation, sight-reading, and rest in professional jazz pianists. By analysing multiple complementary features of the TMS-evoked response — including local field power, phase-locking and maximally reproducible component structure — we aimed to characterize the PMC’s perturbational signature across these distinct performance states. More importantly, we aimed to demonstrate the feasibility of conducting TMS-EEG during a complex and ecologically valid motor task, extending the scope of TMS-EEG research beyond highly simplified motor tasks such as finger tapping.

## Materials & Methods

### Participants

Five right-handed professional jazz pianists (1 female) were recruited for this study. All volunteers were screened for contraindications to TMS (Rossi et al., 2011), including a history of traumatic brain injury, neurological or psychiatric disorders, intracranial metallic implants, drug-release dispensers, metallic tattoos, or cardiac pacemakers. Additional inclusion criteria included: ability to improvise over novel chord progressions on the spot, as depicted in jazz notation; ability to sight-read, as depicted in classical notation; having at least ten years of jazz improvisation experience. Two participants were excluded from the final analysis due to excessive EEG noise. The final sample thus comprised three participants (age: *M* = 30.0, *SD* = 3.5 years; see Table 1).

**Table 1.** Demographic data, musical background and information on experimental sessions for all participants.

| Subject ID | Gender | Age | Formal training (years) | Improvisation experience (years) | Number of jazz concerts | Included? |
| --- | --- | --- | --- | --- | --- | --- |
| S1 | Male | 55 | 40 | 30 | ≈ 700 | No |
| S2 | Male | 33 | 18 | 15 | ≈ 300 | Yes |
| S3 | Male | 26 | 17 | 10 | ≈ 150 | Yes |
| S4 | Male | 30 | 15 | 15 | ≈ 300 | Yes |
| S5 | Female | 25 | 16 | 8 | ≈ 150 | No |

Convenience sampling was used, with recruitment involving outreach to conservatories, university jazz programs, and other relevant institutions. Monetary compensation was not provided. All participants provided written informed consent, and the study was carried out under the ethical standards defined by the Institutional Ethics Committee and the Code of Ethics of the World Medical Association (Declaration of Helsinki).

### Experimental setup overview

This study employed a purpose-designed multimodal TMS-EEG setup tailored for use during live piano performance. The experimental setup integrated four interconnected computers and multiple hardware units (EEG amplifier, TMS and neuronavigation), enabling real-time coordination of stimulation, electrophysiological, and neuronavigational components (Figure 1). Participants performed on a digital piano while neural responses to stimulation and behavioural performance were recorded in parallel. One computer recorded musical performance and generated individualized TMS-click-masking stimuli (Russo et al., 2022), which was mixed with the piano’s audio and delivered via headphones. A second computer acquired EEG data and enabled online monitoring of TMS-evoked potentials (Casarotto et al., 2022). TMS delivery and cross-device synchronization were coordinated by a third computer controlling a central trigger interface. Neuronavigation was managed by a fourth computer, enabling real-time visualization and control of TMS targeting.

**Figure 1.**
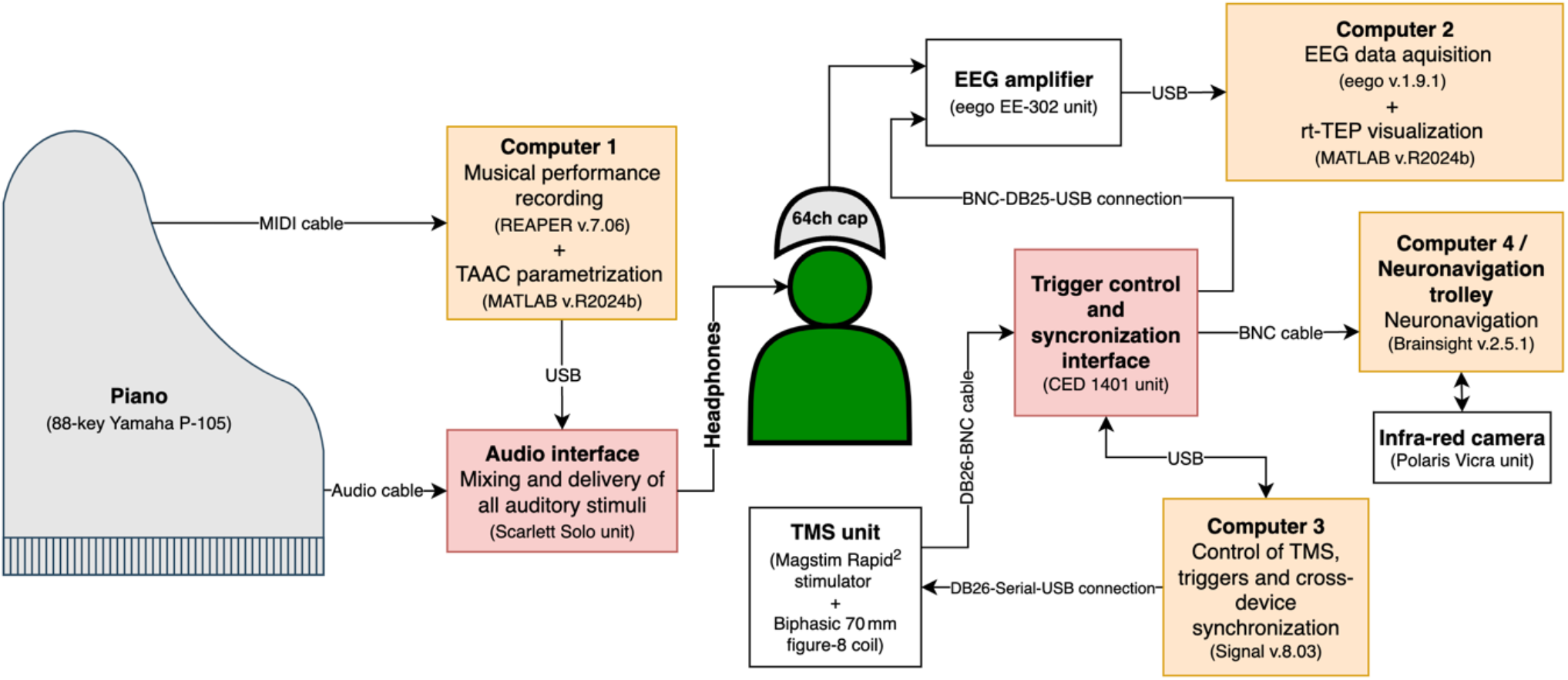
Schematic representation of hardware and software integration.

### TEP recording

TEPs were recorded using a high-density gel-based cap (64-channel cap for TMS with multitrodes; EASYCAP, Germany) comprising sintered Ag/AgCl electrodes with coaxial cabling and arranged in the 10-20 layout with a forehead ground and reference montage. The cap was connected to a TMS-compatible 64-channel amplifier (eego EE-302; ANT Neuro, Netherlands). Impedances were kept below 10 kΩ, signals were band-pass filtered between 0.1–350 Hz and sampled at 8000 Hz with 24-bit resolution. To minimize contamination of TEPs by auditory responses to the TMS coil click (ter Braack et al., 2015), participants wore noise-cancelling in-ear headphones through which individualized masking noise was continuously delivered. This noise was generated using the TAAC toolbox (Russo et al., 2022), which adaptively reproduces the spectral and temporal features of the specific TMS click and superimposes them with calibrated broadband noise to ensure perceptual masking.

### TMS targeting and parametrization

A focal figure-of-eight coil (mean/outer winding diameter ≈ 50/70 mm; biphasic pulse shape; pulse length ≈ 350 µs) connected to a mobile stimulation unit (Magstim Rapid_2_; Magstim Co., UK) was used to deliver single-pulse stimulation. Stimulation targeted the dorsal left premotor cortex (BA 6). A Brainsight 3D infrared Tracking Position Sensor Unit (Rogue Research Inc., Canada) was used to co-register the participant’s scalp anatomy and the TMS coil relative to a standardized template MRI from Brainsight software. Stimulation was delivered within a 2 mm spatial error in relation to the ROI while maintaining angular deviations below 1° for both tilt and orientation.

The identification of artefact-free stimulation targets within the ROI was guided by real-time visualization of TEPs using procedures adapted from the rt-TEP protocol (Casarotto et al., 2022). While this protocol recommends tuning stimulation intensity based on the peak-to-peak amplitude of the earliest TEP component (typically within 0–30 ms), this approach was not feasible due to a prominent recharge artefact that extended up to ≈ 45 ms post-stimulation. As a result, early components were obscured and could not reliably be evaluated. We therefore adapted the procedure by initiating stimulation at each participant’s resting motor threshold and increasing intensity in steps, prioritizing the amplification of the earliest visible component after the artefact window (typically ≥ 45 ms). For each tested intensity, we explored different coil positions, tilts, and orientations within the ROI, using the rt-TEP interface to minimize muscle activity, eye movements, and discharge-related noise. Once no further improvements could be obtained through repositioning, the intensity was increased—unless doing so resulted in excessive artefact duration, in which case the previous setting was retained. The optimal combination of intensity, coil position, and orientation (i.e., the individualized stimulation “hotspot”) was then fixed in the neuronavigation system for the remainder of the session. All subsequent stimulation blocks for that participant — regardless of condition — were delivered using this exact same spatial and stimulation parameterization to ensure consistency.

Stimulation intensity was fixed within each participant and ranged from 60% to 73% of maximum stimulator output (MSO), corresponding to 110% to 115% of their individual resting motor thresholds (RMT). After these preparatory steps, the experimental task began: TMS-evoked responses were recorded in three separate conditions, each comprising at least 200 pulses delivered with a randomly jittered inter-stimulus interval of 2000 to 2300 ms (0.4–0.5 Hz).

### Experimental task

Each participant completed three experimental conditions: jazz improvisation, classical sight-reading, and resting state. All conditions were performed while seated at a digital piano. The first two involved active musical performance, while the third served as a passive baseline. In the improvisation condition (*Improv*), participants were asked to improvise freely on the piano, emulating a solo live performance. Each improvisation was guided by unique 16-bar chord progressions presented in standard jazz notation. These chord sequences were composed specifically for the study, designed to be of comparable difficulty, duration, and structure (4/4 time signature, 120 BPM). In the classical sight-reading condition (*Classical*), participants performed selected pieces from Béla Bartók’s *Mikrokosmos*. These were presented in classical notation and were novel to the participants, who were instructed to perform each piece as accurately and musically as possible at first sight. In the resting state condition (*Rest*), participants remained seated at the piano and were instructed to remain as still and relaxed as possible without engaging in any overt motor or cognitive task.

TEPs were recorded for each condition in separate blocks, and the order of blocks was counterbalanced across participants. The duration of each block varied depending on the time required to deliver at least 200 single TMS pulses. Due to the strict spatial and angular tolerances enforced during stimulation, and the need to pause and readjust the coil when participant movements temporarily disrupted its stability, block durations ranged between 15 and 20 minutes.

### EEG data preprocessing

Data preprocessing was performed on MATLAB R2024b (The MathWorks Inc.) using custom scripts. First, TMS-related artefacts were removed by replacing the signal between –2 and 50 ms relative to the pulse with a clean segment from –9 to –2 ms. This extended window ensured removal of both the pulse artefact and recharge artefact described earlier in the Methods section. Subsequently, trials were segmented from —800 to 800 ms around the stimulus onset and high-pass filtered at 1 Hz. Trials and channels were visually inspected and manually rejected when exhibiting excessive noise. Channels were then re-referenced to the average reference, after which independent component analysis (ICA) was applied to identify and remove residual artefacts related to eye movements, scalp muscle activity, and device interference (EEGLAB runica function; Delorme & Makeig, 2004). Interpolation of rejected channels was performed after ICA, followed by downsampling to 1 kHz.

### TEP analysis

Analyses were implemented in MATLAB using custom code operating on EEGLAB-format datasets (EEG structs). We compared *Improv, Classical* and *Rest* in either single-participant or group mode. Unless noted, statistics are nonparametric and family-wise error rate (FWER) controlled via permutation tests (*α* = .05).

#### Divergence Index (DI)

DI provided a preliminary global, time-resolved estimate of how much TMS-evoked EEG responses differed pairwise between conditions by quantifying the proportion of post-stimulus samples showing statistically significant amplitude differences across all channels (Casarotto et al., 2010). After baseline handling via iterative Wilcoxon-based baseline matching (*α* = .05), condition means were contrasted sample-by-sample from 50–250 ms post-stimulus. Signals were band-pass filtered at 1–45 Hz and baseline-subtracted using −250–0 ms. FWER across time was controlled with 10,000 label-shuffle permutations (time-wise max-statistic). The DI value is the percentage of post-stimulus samples declared significant. The empirical cutoff of 1.7% (Casarotto et al., 2010) was used as a reference threshold. As an additional control, we computed a subject-specific split-half baseline distribution for Rest (1,000 random half-splits; 1,000 permutations for the threshold), providing a within-subject reference for expected DI under no systematic between-condition differences.

#### Local mean field power (LMFP)

LMFP assessed the strength of TMS-elicited currents over the four electrodes directly beneath the coil (FC1, FCz, C1, Cz) (a proxy for local cortical excitability; Fecchio et al., 2017). Per-trial signals were band-pass filtered at 1–45 Hz, baseline-corrected with the mean over −400 to −50 ms and transformed as the root-mean-square across the four ROI channels at each time point. Values were summarized in 50–300 ms. Within-condition enhancement relative to baseline was tested via one-sided sign-flip permutations across trials with family-wise error control over time using the maximum statistic (*α* = .05; 5,000 permutations). Between-condition differences were tested with label-shuffle permutations (FWER over time via max-statistic; 5,000 permutations).

#### Broadband phase-locking factor (bPLF)

To quantify phase consistency of the evoked response, we computed PLF over the same 4-channel ROI in a broadband 8–45 Hz band. Trials were band-pass filtered (4th-order Butterworth), analytic phase was obtained with the Hilbert transform, PLF was computed per channel and averaged across the ROI, and statistics were restricted to 50–300 ms. Within-condition significance was assessed with phase-randomization permutations: on each permutation, every trial’s phase time series was offset by a random constant chosen uniformly from zero to one full cycle (0–360°), PLF was recomputed, and the maximum PLF within 50–300 ms served as the test statistic to control family-wise error over time (*α* = .05; 5,000 permutations). Between-condition differences were tested with label-shuffle permutations at the trial level, using the max absolute PLF difference within 50–300 ms as the statistic (FWER by max-statistic; 5,000 permutations).

#### Correlated Components Analysis (CorrCA)

We used CorrCA to test whether TMS evokes common, reproducible network-level responses within and across conditions (Parra et al., 2019). CorrCA identifies linear combinations of EEG channels whose time courses are maximally correlated across trials within a post-stimulus window. Mathematically, CorrCA solves a generalized eigendecomposition that maximizes the ratio of between to within-trial covariance, which is equivalent to maximizing inter-trial correlation. CorrCA was run on the four under-coil electrodes (FC1, FCz, C1, Cz) in a 50–150 ms window. Trials were entered into a regularized CorrCA, with the shrinkage parameter (*γ*) chosen by split-half cross-validation. Significance of inter-trial correlations (ISCs) was assessed with circular time-shift permutations (10,000 surrogates) using trimming-aware nulls and family-wise error control by the component-wise max statistic (*α* =.05). “Transfer” was evaluated by applying filters learned in a reference condition to target conditions and testing the resulting ISCs against each target’s permutation distribution; evidence is reported as −log_10_(*p*_max_).

Cross-condition correspondence was quantified two ways. First, we matched components one-to-one by absolute map correlation (|*r*_map_|) and compared their ERPs via Pearson correlation after sign-alignment. We report *r*_ERP_at zero lag and with a small-lag search (± 10 ms). Second, we did strict C1 to C1 comparisons: (i) |*r*_map_| for C1 forward maps, (ii) *r*_ERP_with max-lag (± 10 ms) after the same sign rule, (iii) the angle between C1 weight vectors, and (iv) a confirmatory waveform check after an orthogonal rotation within the shared subspace (L = 2). For figures, component traces are shown as mean ± SEM with a display-only demean on 48–52 ms; plotting normalization does not affect statistics. Transfer evidence for C1 was summarized in both directions (A to B and B to A) and we report the conservative single value as the minimum of the two −log_10_(*p*_max_) estimates.

## Results

### Divergence Index

DI values revealed clear differences in TMS-evoked EEG responses across conditions and participants (Figure 2). All pairwise contrasts showed DI values that substantially exceeded each participant’s split-half baseline threshold and the empirical cutoff of 1.7% (Casarotto et al., 2010), suggesting that the observed differences reflect systematic condition effects rather than intrinsic variability.

**Figure 2.**
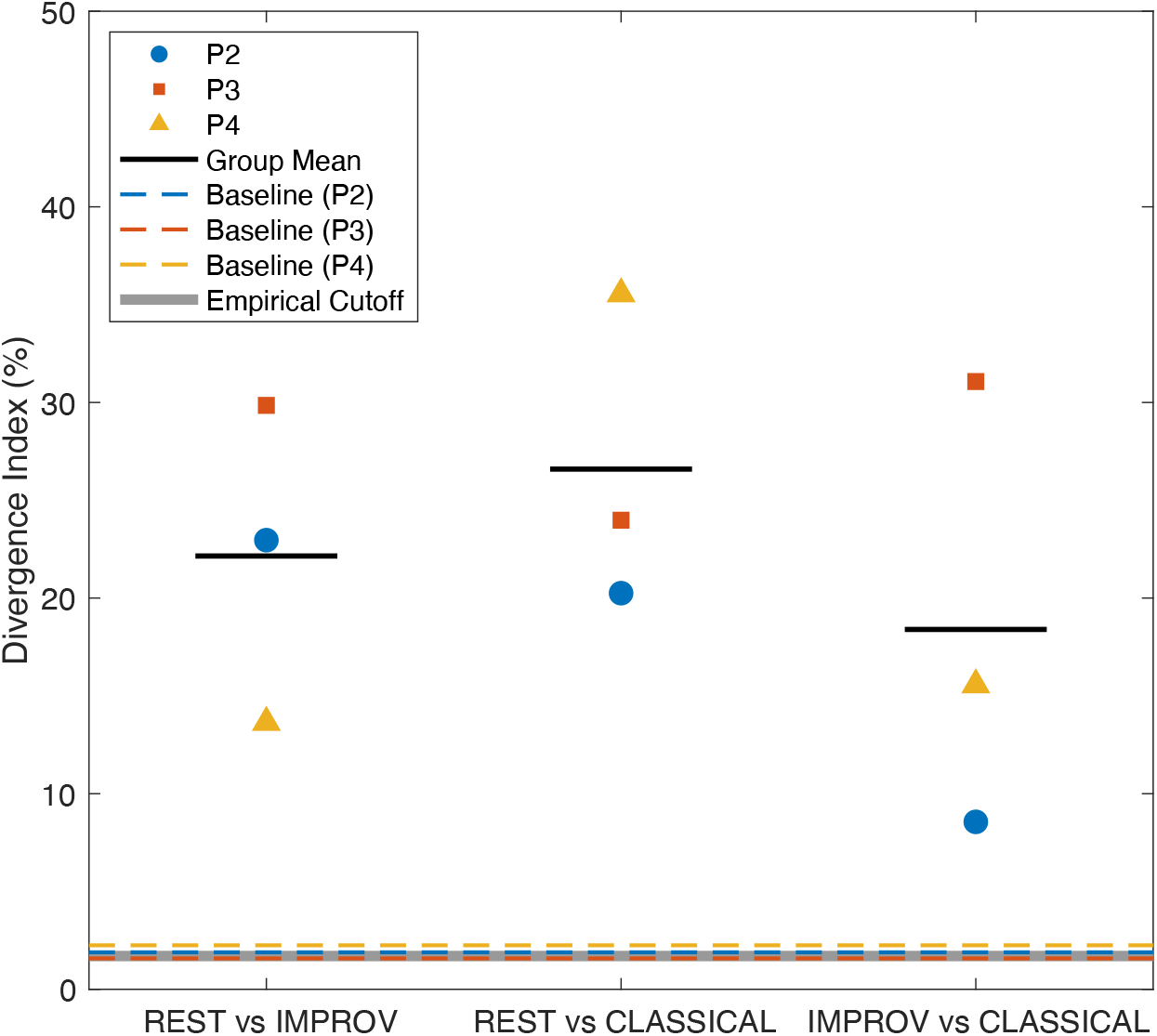
Divergence index by condition contrast and participant.

### Local mean field power (LMFP)

LMFP was quantified over the four electrodes directly beneath the coil (FC1, FCz, C1, Cz) across the 50–300 ms analysis window (Figure 3). Overall power was highest for *Rest* (*M* = 0.77 µV) and *Classical* (*M* = 0.76 µV), and markedly lower for *Improv* (*M* = 0.45 µV). Significant between-condition contrasts (two-sided FWER < .05), visualised as horizontal black bars beneath the LMFP traces, revealed divergence for *Rest* vs *Improv* (41.3% of the window) and *Improv* vs *Classical* (31.0%), whereas *Rest* vs *Classical* showed no reliable differences. Permutation tests confirmed that these responses were not baseline noise. Within-condition comparisons showed that 98.4% of *Rest* samples, 86.5% of *Classical* samples, and 36.5% of *Improv* samples exceeded their family-wise thresholds (one-sided FWER < .05). Thus, *Rest* and *Classical* elicited similarly strong cortical currents, while *Improvisation* was consistently weaker, driving all significant pairwise effects highlighted in the figure

**Figure 3.**
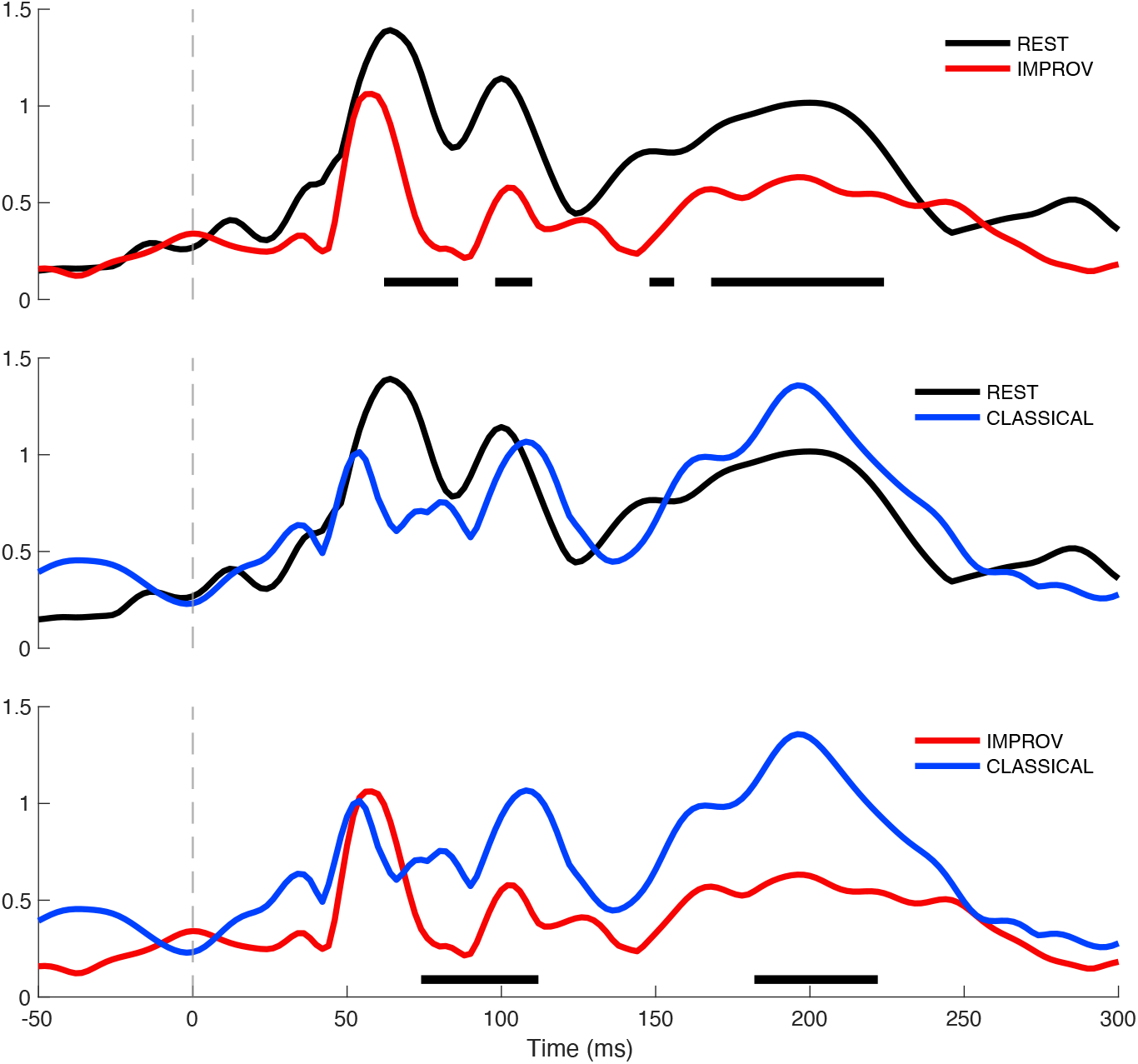
Local mean field power across conditions Note. The three panels show pairwise contrasts between conditions. Horizontal black bars beneath the traces denote time samples with significant differences between the two conditions in each panel.

### Local broadband phase-locking factor (bPLF)

A broadband (8-45 Hz) PLF was quantified over the four electrodes directly beneath the coil (FC1, FCz, C1, Cz) across the 50–300 ms analysis window. All three conditions exhibited significant broadband phase alignment within 50–300 ms, but to markedly different degrees (see Figure 4). *Rest* showed the most sustained locking, with 43.7% of samples exceeding the permutation threshold (threshold = .097), a peak bPLF of .301, a 110 ms contiguous supra-threshold segment, and the largest area-under-the-curve (AUC = 7.33). *Classical* showed a similar profile: 34.1% significant (threshold = .102), the highest peak amplitude (.364), a 26 ms duration, and AUC = 7.77, whereas *Improv* yielded the weakest and least sustained locking (15.9% significant, threshold = .102; peak = .246; duration = 24 ms; AUC = 2.41). Pairwise permutation tests revealed limited inter-condition effects: *Rest* differed from *Improv* on 9.5% of the window (|threshold| = .101), *Rest* and *Classical* did not differ (0%), and *Improvisation* differed from *Classical* on just 4.8% of samples (|threshold| = .116). Overall, bPLF results indicate stronger and more sustained stimulus-locked phase synchrony during *Rest* and *Classical* than during *Improvisation*, with minimal differences between former two.

**Figure 4.**
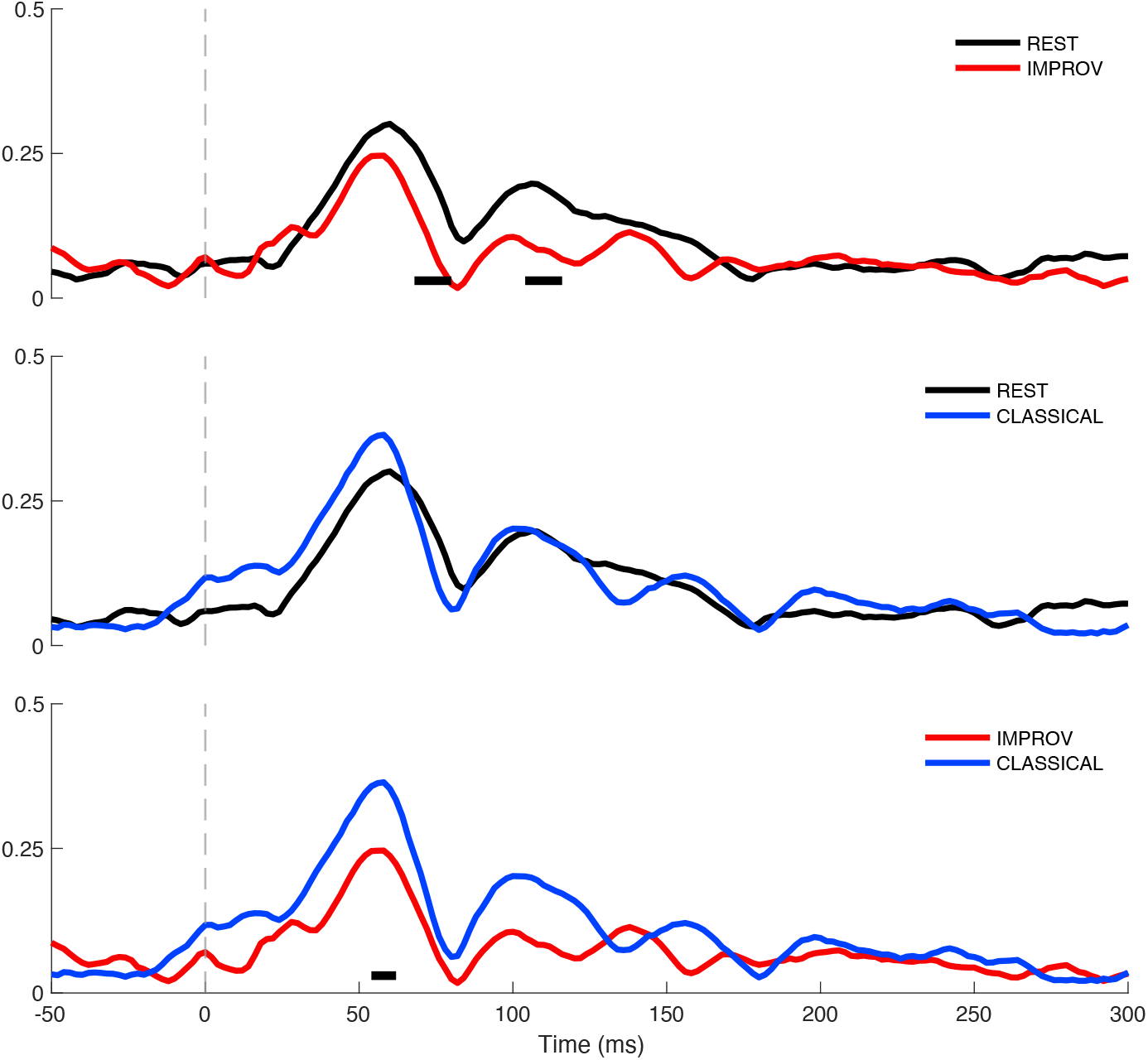
Local broadband phase-locking factor across conditions. *Note*. The three panels show pairwise contrasts between conditions. Horizontal black bars beneath the traces denote time samples with significant differences between the two conditions in each panel.

### Correlated Components Analysis (CorrCA)

We applied CorrCA over the four electrodes directly beneath the coil (FC1, FCz, C1, Cz) across a focused 50– 150 ms analysis window to avoid later sensory components. CorrCA recovered reliable components in all conditions, with markedly stronger within-condition structure in *Rest* and *Classical* than in *Improv*. The leading inter-trial correlation was comparable for *Rest* (ISC_1_ ≈ .106) and *Classical* (ISC_1_ ≈ .106), but smaller for *Improv* (ISC_1_ ≈ .060). Permutation testing confirmed multiple significant components (max-statistic: *Rest* = 3, *Classical* = 2, *Improv* = 1). A *γ* sweep showed ISC_1_ was stable to regularization, indicating these effects are not driven by the shrinkage choice. Cross-condition transfer for C1 was at the permutation floor for 10,000 permutations (−log_10_(*p*_max_) = 4.00) for all pairings, indicating very strong generalization. For visualization of Component 1 (C1), baseline-corrected overlays are shown in Figure 5a, and raw-amplitude overlays under a fixed *Rest* projection (*W*_ref_) are shown in Figure 5b.

**Figure 5.**
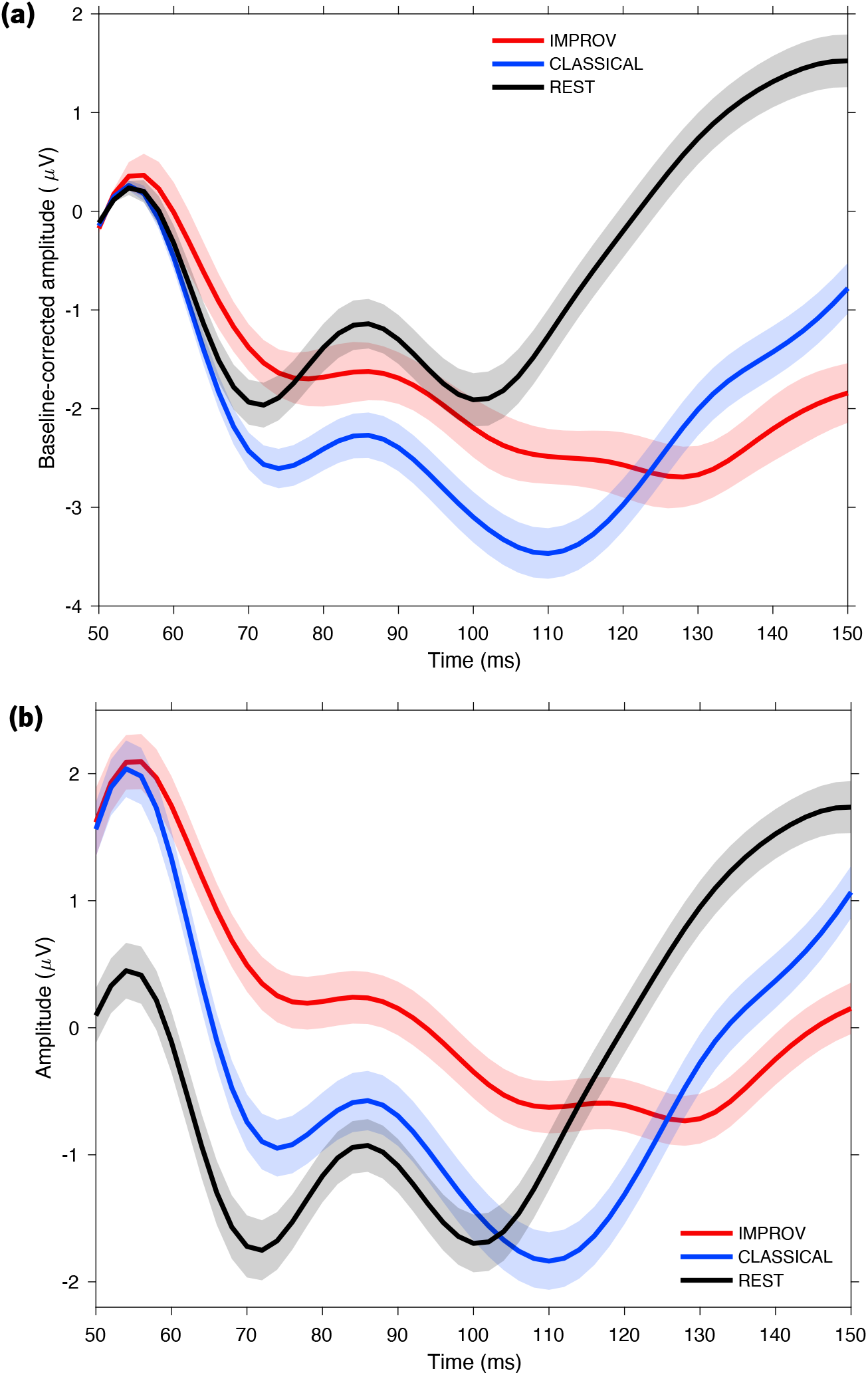
Component 1 (shape-first) — baseline-corrected overlays. *(b)* Component 1 (amplitude-first) — overlays under fixed Rest projection. *Note*. (a) Each trace represents CorrCA Component 1 per condition, demeaned per trial on 48–52 ms and polarity-aligned to emphasize waveform morphology across conditions, independent of overall amplitude. Displays mean ± SEM in µV. (b) All conditions are projected with the Rest spatial filter (*W*_ref_) and shown in raw µV with polarity aligned. This display highlights between-condition amplitude differences while preserving a comparable time course.

Using *Rest* as reference, cross-condition matching by forward maps yielded strong topographic correspondence for C1 with *Classical* (|*r*_map_| = 0.97) and substantial correspondence with *Improv* (|*r*_map_| = 0.86). ERP similarity at ± 10 ms was high for *Classical* (*r* ≈ 0.62) and moderate for *Improv* (*r* ≈ 0.37) in the shape-first view (Figure 5a), but when projecting all conditions with *W*_ref_the C1, ERPs converged for both targets (*r* ≈ 0.59–0.64), and a simple L = 2 subspace alignment further equalized them (*r* ≈ 0.62; values summarized in Table 2). Consistent with this, C1 transfer remained at the permutation floor (−log_10_(*p*_max_) = 4; see Figure 6, “Transfer”). Principal-angle analyses further support a rotation account: *Classical*’s first weight dimension is nearly parallel to Rest (*θ*_1_ ≈ 1.8°), whereas Improv shares the first dimension more modestly (*θ*_1_ ≈ 16.7°) and diverges in the second (*θ*_2_ ≈ 62.8°), consistent with a low-dimensional rotation rather than a wholesale change.

**Table 2.**
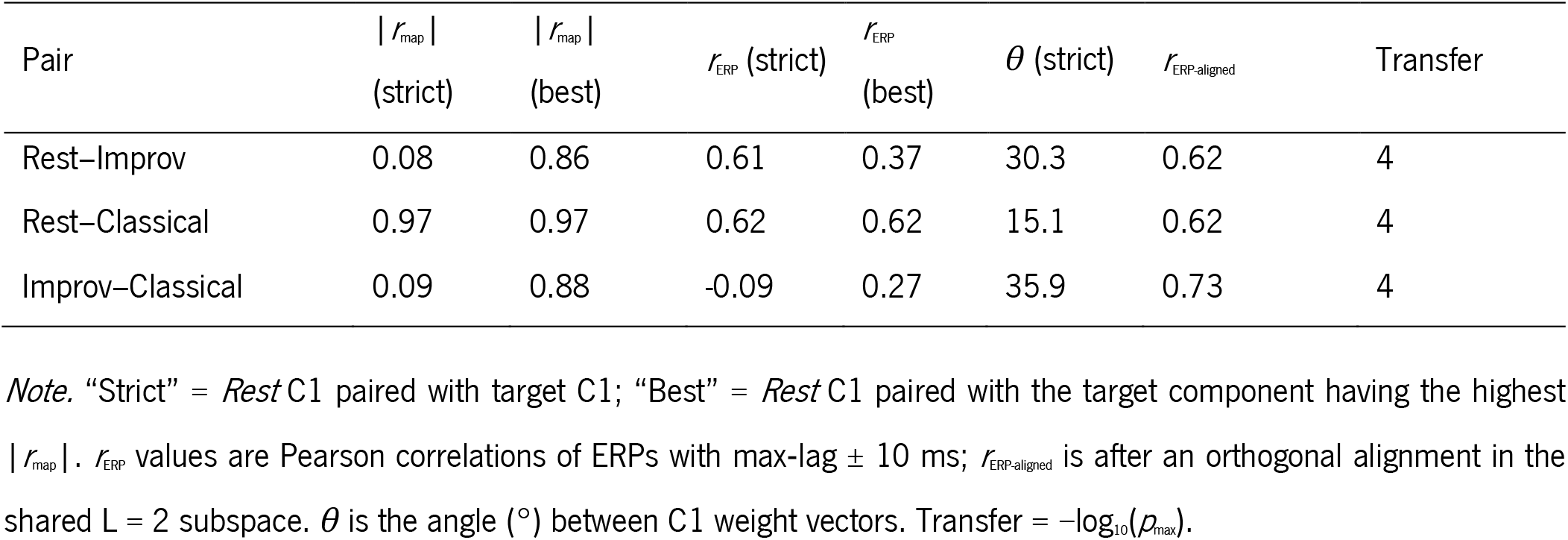
Summary of CorrCA C1 similarity and generalization across conditions.

**Figure 6.**
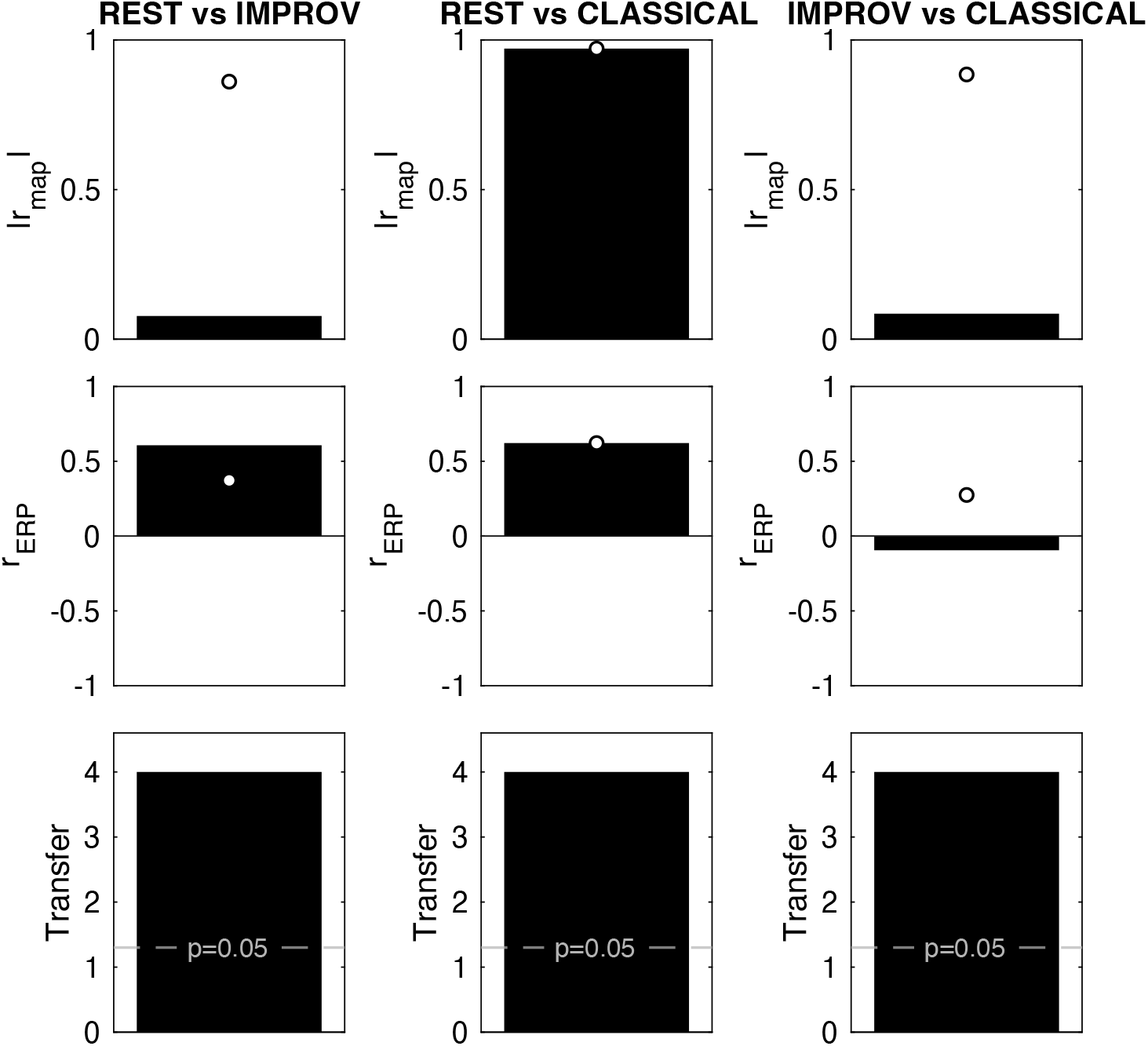
Component 1 similarity summary (pairwise) *Note*. Bars summarize (from top to bottom): |*r*_map_| between forward maps, *r*_ERP_(max-lag, +/-10 ms), and transfer evidence reported as −log_10_(*p*_max_) using its minimum value. White circles mark best-match overlays (C1 paired to the target’s most similar component by |*r*_map_|). Metrics not shown as bars (angle and aligned *r*_ERP_) are reported in Table 2.

A strict pairwise check that fixes the index to “C1 vs C1” highlights this rotation and is summarized in the bar panels (Figure 6): *Rest*–*Improv* shows a very low C1-to-C1 map correlation (|*r*_map_| = 0.08) even though ERP similarity (*r* ≈ 0.61) and bidirectional transfer evidence (−log_10_(*p*_max_) = 4) remain strong; *Rest*–*Classical* shows high agreement on all metrics (|*r*_map_| = 0.97; *r* ≈ 0.62; −log_10_(*p*_max_) = 4). After within-subspace alignment, the ERP similarity for the “C1 process” recovers to high values (*r* ≈ 0.62–0.73) even for pairs involving *Improv* (Table 2), indicating that *Improv*’s C1 is effectively a rotated/reindexed combination of the first few *Rest* components. In short, the dominant response around 50–150 ms is preserved across conditions. Discrepancies with *Improv* in strict C1-to-C1 map comparisons reflect component mixing within the shared subspace, not a polarity issue or a loss of the underlying effect.

## Discussion

This study employed TMS-EEG to investigate state-dependent dynamics of the left dorsal premotor cortex during musical improvisation, sight-reading, and rest in professional jazz pianists. Our findings reveal distinct perturbational signatures that differentiate improvised performance from both sight-reading and resting states, providing novel causal insights into how the PMC dynamically reconfigures its response properties to support different modes of motor control.

### Reduced local excitability during improvisation

Our most striking finding was the marked reduction in LMFP during improvisation compared to both rest and classical sight-reading conditions. The approximate 41% reduction relative to rest and classical performance, with no significant difference between rest and classical conditions, suggests that improvisation engages the PMC in a fundamentally distinct excitability state, one in which TMS perturbation elicits substantially smaller local activation.

This pattern aligns with a state-dependent gating mechanism whereby improvisation places PMd in a mode that selectively attenuates responses to exogenous, time-locked stimulation while maintaining ongoing, internally guided computations. Previous TMS-EEG studies have demonstrated similar dampening of evoked potentials during active motor states compared to rest, including reduced N100 and late positive components over motor areas during simple movement execution or inhibition (Nikulin et al., 2003; Yamanaka et al., 2013), consistent with decreased susceptibility of local circuits when they are behaviourally engaged. Our findings extend this principle to complex creative motor behaviour, suggesting that the task-dependent reduction in cortical responsiveness scales with the cognitive demands and internal focus required for improvisation.

The reduced LMFP during improvisation converges with neuroimaging evidence supporting a neural efficiency account of musical improvisation expertise. Expert improvisers show paradoxically lower BOLD activation across frontoparietal executive regions alongside stronger functional connectivity between bilateral DLPFC, PMd, and pre-SMA (Pinho et al., 2014, 2015). This pattern has been interpreted as reflecting the automation and streamlining of creative motor operations through optimized premotor-prefrontal coupling (Beaty, 2015; Beaty et al., 2016). This finding also aligns with Haslinger and colleagues’ (2004) finding of reduced activation of PMd (among other motor network regions) in concert pianists during bimanual coordination, which was argued to be indicative of a training-induced efficiency of cortical systems that could be fundamental to achieve high-level motor skills.

### Preserved early components with reduced gain

Despite the overall reduction in evoked power, CorrCA (which maximizes inter-trial correlation, equivalent to maximizing repeat-reliability of the evoked waveform; Parra et al., 2019) revealed that the fundamental structure of the early cortical response (50-150 ms) remains preserved across all conditions. The dominant component showed strong generalization between rest and classical conditions (|*r*_map_| = 0.97) and substantial correspondence even with improvisation after accounting for subspace rotation. This dissociation (i.e., preserved component morphology with reduced amplitude) suggests that improvisation does not fundamentally alter the canonical premotor response architecture but rather modulates its gain.

This finding reconciles heterogeneity in the literature regarding premotor involvement in improvisation. While some studies report increased PMd activation during improvisation (de Manzano & Ullén, 2012), others show decreased activity with expertise (Pinho et al., 2014). Our results suggest that both observations may reflect different aspects of the same phenomenon: the PMd remains engaged in the same fundamental computations (preserved component structure) but operates in a more efficient, less excitable regime (reduced LMFP). This interpretation aligns with theories proposing that expertise involves not the recruitment of different neural circuits but rather the optimization of existing ones (Beaty, 2015; Beaty et al., 2016; Pinho et al., 2014).

### Reduced broadband phase-locking

This reduced PLF indicates that while TMS still evokes a reproducible response during improvisation (as shown by CorrCA), the timing of this response is more variable across trials. During rest and classical sight-reading, the PMd maintains more stereotyped response timing to external perturbation, with each TMS pulse triggering a response at a consistent phase. In contrast, during improvisation, the temporal relationship between stimulation and response becomes less deterministic.

This finding aligns with the reduced LMFP in suggesting that improvisation engages a cortical state less dominated by stimulus-locked responses. The variable response timing during improvisation may indicate that ongoing endogenous dynamics play a stronger role in shaping when and how the cortex responds to perturbation. This could reflect: (i) stronger ongoing oscillations that interact differently with the TMS pulse on each trial, (ii) a more complex state space where perturbations can follow multiple trajectories, or (iii) active gating mechanisms that modulate response timing based on the current phase of internal processes. Together with our other local measures, this temporal variability further characterizes the distinct cortical state associated with improvisation, one that maintains responsiveness (preserved components) but with reduced amplitude (LMFP) and less predictable timing (PLF).

Because inter-trial phase metrics can be influenced by differences in evoked amplitude and latency variability (van Diepen & Mazaheri, 2018), we interpret the PLF attenuation with appropriate caution. Nevertheless, the use of phase-randomization and family-wise error statistical controls, together with the concordant LMFP decrease in the same ROI, makes a purely power-driven account unlikely.

### Limitations and future directions

Several limitations warrant consideration. First, our sample size of three participants limits the generalizability of our findings. While the high internal validity of TMS-EEG studies, arising from the large number of observations per participant per condition, allows us to establish effects with near certainty within our sample, extending these findings to the broader population of jazz musicians warrants caution. The demanding nature of these experimental sessions, which lasted up to five hours and required participants to perform complex musical tasks while maintaining precise coil positioning, made larger-scale data collection exceptionally challenging. This study thus represents a proof-of-concept and methodological innovation rather than a systematic investigation, demonstrating the feasibility of combining TMS-EEG with real musical performance despite the considerable technical and practical obstacles.

Second, the presence of recharge artefacts limited our analysis to responses beyond 50 ms. Current-generation TMS systems with customisable coil recharge timing would eliminate this limitation, allowing future investigations to probe these earlier components. Additionally, while our focus on the left PMd was motivated by its established role in internally generated movement, improvisation likely involves distributed networks extending beyond this single region. Future studies employing multiple stimulation sites could map how different nodes of the motor network reconfigure during creative performance.

## Conclusions

This study provides initial causal evidence that musical improvisation fundamentally alters the response properties of the premotor cortex to external perturbation. The convergent findings of reduced local excitability, preserved but gain-modulated components, and decreased phase-locking support a neural efficiency model in which improvisation engages a distinct cortical state optimized for internally driven motor control. More broadly, our findings demonstrate the value of combining perturbational approaches with ecologically valid tasks to understand the neural basis of complex human abilities. By directly probing how cortical circuits respond to external perturbation during real musical performance, we reveal state-dependent dynamics that would remain hidden in traditional activation or correlational studies. This approach opens new avenues for understanding how the brain flexibly reconfigures its computational properties to support the human capacity for creative expression.

## Research transparency statement

### Data availability

Datasets, analysis scripts and materials supporting this study will be made available upon publication.

### Ethics approval statement

This study was approved by the Ethics Committee of Faculty of Psychology and Educational Sciences, University of Coimbra (reference code: CEDI/FPCEUC:64/3).

